# Landscape of microRNA and target expression variation and covariation in single mouse embryonic stem cells

**DOI:** 10.1101/2024.03.24.586475

**Authors:** Marcel Tarbier, Sebastian D. Mackowiak, Vaishnovi Sekar, Franziska Bonath, Etka Yapar, Bastian Fromm, Omid R. Faridani, Inna Biryukova, Marc R. Friedländer

## Abstract

MicroRNAs are small RNA molecules that can repress the expression of protein coding genes post-transcriptionally. Previous studies have shown that microRNAs can also have alternative functions including target noise buffering and co-expression, but these observations have been limited to a few microRNAs. Here we systematically study microRNA alternative functions in mouse embryonic stem cells, by genetically deleting *Drosha* - leading to global loss of microRNAs. We apply complementary single-cell RNA-seq methods to study the variation of the targets and the microRNAs themselves, and transcriptional inhibition to measure target half-lives. We find that microRNAs form four distinct co-expression groups across single cells. In particular the mir-290 and the mir-182 clusters are abundantly, variably and inversely expressed. Intriguingly, some cells have global biases towards specific miRNAs originating from either end of the hairpin precursor, suggesting the presence of unknown regulatory cofactors. We find that miRNAs generally increase variation and covariation of their targets at the RNA level, but we also find miRNAs such as miR-182 that appear to have opposite functions. In particular, miRNAs that are themselves variable in expression, such as miR-291a, are more likely to induce covariations. In summary, we apply genetic perturbation and multi-omics to give the first global picture of microRNA dynamics at the single cell level.

## Background

MicroRNAs (miRNAs) are ∼22 nucleotide small non-coding RNAs that guide Argonaute effector proteins to target mRNAs in a sequence-specific manner, leading to their translational inhibition and increased degradation through de-adenylation and de-capping (Bartel 2018). miRNAs group into seed families that mostly target the same pool of mRNAs through complementary base pairing of their seven-nucleotide seed region (Lai 2002; Bartel 2009). Each family targets hundreds of mRNAs and together miRNAs are considered key post-transcriptional regulators (Friedman et al. 2009).

It is well-established that miRNAs can repress the expression of their targets, both at the RNA and the protein level (Baek et al. 2008; Selbach et al. 2008). However, alternative functions for miRNAs have also been proposed, including buffering of *gene expression variation* (noise) (Hornstein and Shomron 2006; Tsang et al. 2007; Ebert and Sharp 2012; Siciliano et al. 2013). This is an attractive hypothesis, since miRNAs reduce translational efficiency of their targets, which reduces the noise of their targets. Similarly, the repression allows higher transcription of the targets to reach the same final protein output, and this higher transcription again reduces noise according to theory. The hypothesis that miRNAs buffer gene expression noise fits well with observations that many mutant animals that are devoid of specific miRNAs do not display obvious phenotypes under laboratory conditions, but only when subjected to stress conditions (Miska et al. 2007; Li et al. 2009).

Another alternative miRNA function that has been proposed is to induce *gene expression covariances* between targets (synchronize expression) (Rzepiela et al. 2018; Tarbier et al. 2020). In individual cells where a given miRNA is highly abundant, the targets will be coordinately repressed, while in a cell where the miRNA is lowly abundant, the targets will be coordinately alleviated. This would for instance make sense for targets that form part of the same protein complex, where correct stoichiometry is important for correct assembly and folding (Tarbier et al. 2020; Gutierrez-Perez et al. 2021). However, studying these alternative functions require single-cell methods, since variation and covariation across individual cells must be measured.

There is evidence for both of these alternative miRNA functions. In 2015, a study combining fluorescent reporters bearing binding sites for select miRNAs with single-cell imaging showed that miRNAs can reduce target expression noise for lowly abundant targets, and increase target expression noise for highly abundant targets, at the protein level(Schmiedel et al. 2015). This could have the biological function of ensuring that the exactly right amount of protein is output in each single cell, which can be important for lowly abundant regulators like transcription factors.

Since then, reliable methods to measure both miRNAs and their targets in single cells using RNA sequencing have emerged. Standard single-cell RNA-sequencing methods such as 10x Chromium or Smart-seq3 can detect mRNA targets with high throughout in terms of cells, or high sensitivity in detecting individual mRNA molecules, respectively (Zheng et al. 2017; Hagemann-Jensen et al. 2020). There also exists several protocols to sequence miRNAs in single cells, which are quite similar to the methods used for pooled cell (bulk) applications, but with minor modifications. These methods give representative views of the miRNA profiles of single cells, although they typically detect only a subset of the miRNA molecules in the samples(Faridani et al. 2016; Hucker et al. 2021).

A recent study applied an innovative but labor-intensive method to split the lysate from single cells in two, and sequence the small RNA complement from one half and the mRNA complement from another half (Wang et al. 2019). This study provided evidence that miRNAs are negatively correlated with their targets in single cells, and that miRNAs themselves can vary in expression between cells. Another study provided evidence that miR-182 can cause its targets (including several known pluripotency factors such as Nanog and Sox2) to vary more in expression, and that this increased variation is important for transitions between mouse embryonic stem cell equilibrium states (Chakraborty et al. 2020). Two independent studies introduced a total of four miRNAs into cells and observed that their targets increased in variation but also that the targets started to covary more (Gambardella et al. 2017; Rzepiela et al. 2018). This covariation was also observed in recent study from our group, where we observed expression covariations between targets disappear in mouse embryonic stem cells that were genetically depleted of the key Drosha biogenesis protein (Tarbier et al. 2020).

In summary, there is emerging evidence that miRNAs may have other function than simple repression of their targets, including increasing or buffering target expression variation at the RNA or protein level. However, previous studies have focused on limited numbers of miRNAs and targets, and no systematic transcriptome-wide investigation has yet been undertaken.

Here we profile the biogenesis and function of miRNAs in single embryonic stem cells by globally profiling miRNAs in single cells, mRNA transcripts in single cells and mRNA half-lives in the same cell population. We apply a previously published method - Small-seq (Faridani et al. 2016) - to sequence miRNAs in 192 individual mouse embryonic stem cells, more cells than have been profiled in previous miRNA single-cell studies. We find that specific miRNA families are highly variable across cells, even in our homogenous cell population. Integrating our Small-seq data with public measurements of miRNA half-life, we find that short-lived miRNAs tend to be more stably expressed. We study miRNA covariation patterns and find that miRNAs expressed from the same genomic clusters tend to be positively correlated, suggesting co-transcriptional regulation that is preserved to the level of the mature miRNAs. We find that miRNAs from the mir-290 cluster and the mir-182 clusters are strongly and negatively correlated across single cells. Since these miRNAs are known to have overlapping functions in pluripotency but distinct functions in regulation of cell cycle and cell transitions (see Discussion section), this suggests the presence of a miRNA-mediated functional switch. We find that some individual cells have global biases towards miRNAs originating from either the 5-prime arm or 3-prime arm of the precursor transcript, suggesting the existence of currently unknown cofactors that determine this bias.

Further integrating our previously published whole-cell single-cell RNA sequencing data, we find that miRNA primary transcripts are abundantly detected, as cleaved transcripts in control mouse embryonic stem cells and as full-length transcripts in cells that are genetically ablated for the Drosha biogenesis protein. We find that miRNA-mediated repression is relatively low (∼15% repression) compared to the overall natural variation of target expression between cells. Expanding findings from other studies, we find that most miRNAs induce target variation and covariation of their targets at the RNA level, but we also find notable examples of miRNAs with opposite effects. Finally, we show that the induction of gene expression covariation in the target pools of miRNAs can be directly linked to the expression variability of the miRNAs themselves. In summary, we here combine Small-seq with single-cell mRNA sequencing to give a more detailed look into miRNA functions at the cellular level, supporting and extending previous observations, and giving new insights into miRNA biogenesis and covariance patterns.

## Results

### Small-seq detects ∼3000 distinct miRNA molecules in each mouse embryonic stem cell

We sequenced the miRNA complements of 192 individual mouse embryonic stem cells using an optimized Small-seq (single-cell small RNA-seq) (Faridani et al. 2016) protocol to investigate miRNA variability in these cells. The cells had been cultured in a 2i and LIF medium and sorted in the G2/M cell cycle stage to generate a homogenous population (Methods)(Kolodziejczyk et al. 2015). Following stringent quality control (Suppl. Fig. 1-2) we found that the number of detected distinct miRNA sequences ranged from 160 to 300, with an average of 230 distinct miRNAs (Figure 1A, Supplementary Table S1). This number is higher than that reported in a recent benchmarking study (Hucker et al. 2021), highlighting the high quality of our data. We found that the miRNA with the highest expression was miR-290a-5p, which was detected in ∼500 molecules per cell on average (Figure 1B). Summing over all miRNAs, we found an average of ∼3000 molecules per cell. Assuming that one mouse embryonic stem cells contain ∼110,000 miRNA molecules (Calabrese et al. 2007) (Calabrese et al., 2007), comparable to measurements in other mammalian cells (Bissels et al. 2009), this suggests that the sensitivity of our method is around ∼3%. We found that the miRNA composition is relatively similar across cells, showing mostly subtle differences (Figure 1C). On average, the ten most highly expressed miRNAs comprised 55% of the total miRNA molecules in a given cell, comparable to bulk data (Lappalainen et al. 2013). In summary, we find that miRNA complements in single mouse embryonic stem cells can be profiled to fine resolution, even though the sequencing has relatively low sensitivity.

**Figure 1:**
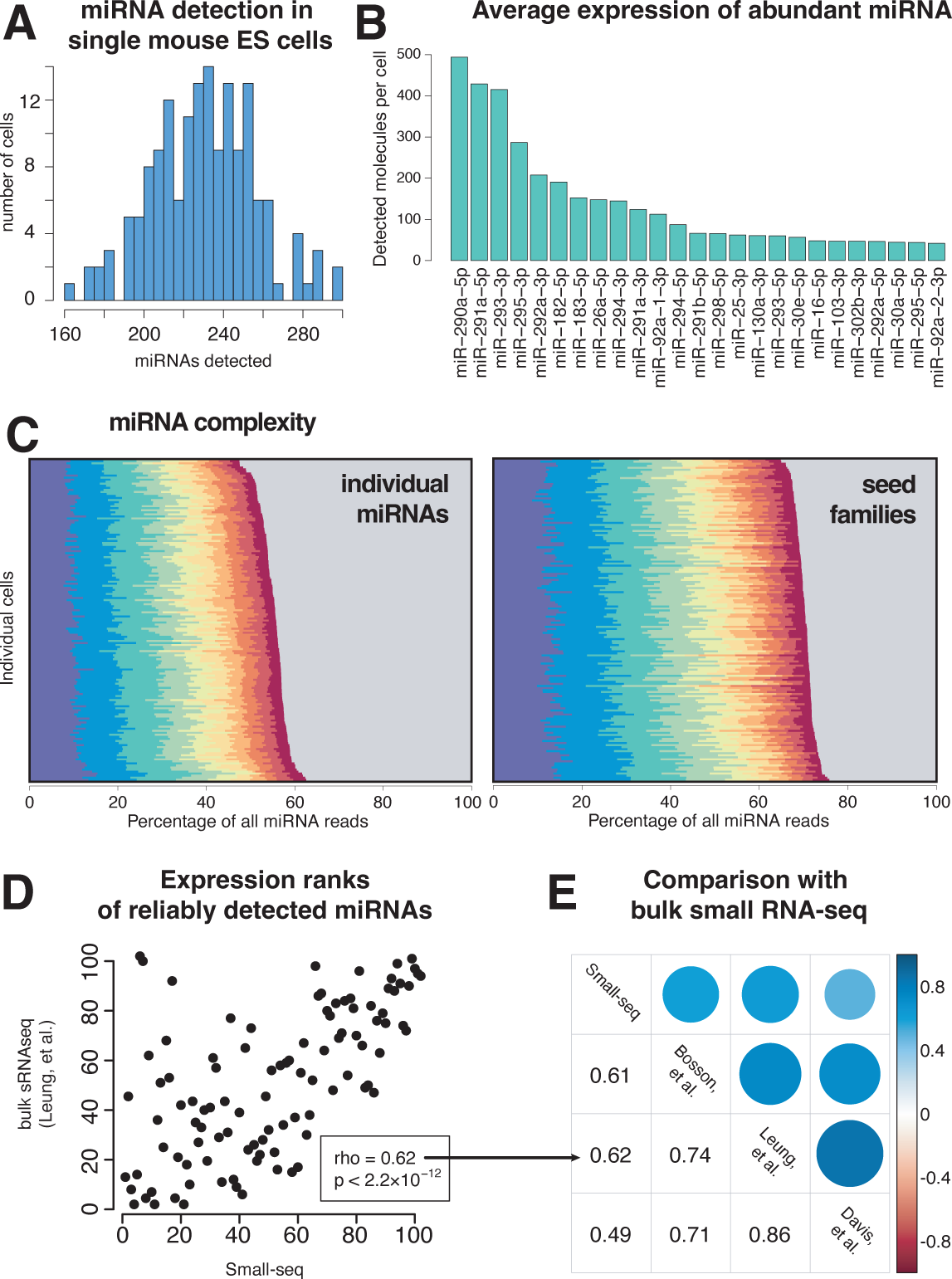
miRNA profiling in single mouse embryonic stem cells by Small-seq. **A** Number of distinct miRNAs detected in each single cell. **(B)** Top 25 miRNAs in mouse embryonic stem cells. The number of detected molecules was estimated using unique molecular identifiers (UMIs) to resolve PCR duplicates. **(C)** Expression of top 10 miRNAs in 192 individual single cells. The grey area represents the contribution of all miRNAs other than the top ten ones. **(D)** Expression rank of miRNAs detecting in mouse embryonic stem cells by Small-seq and bulk small RNA-seq (Leung 2011). Lower rank means higher expression. **(E)** Spearman’s rank correlation between miRNA expression as profiled by Small-seq and three representative bulk small RNA-seq data sets (Leung 2011, Bosson 2014, Davis 2012).

### Small-seq data correlates well with bulk small RNA-seq data

The miRNA complement of mouse embryonic stem cells have been profiled in bulk cells in several studies. We find that the expression ranks of miRNAs in bulk small RNA-seq data correlates well with Small-seq (here the data from all single cells were pooled, Figure 1D). The Spearman’s correlation coefficient of miRNA expression ranks ranged from 0.49 to 0.61 when compared with three representative bulk studies (1E). Some differences may in fact reflect biological differences in culture conditions—all available bulk sRNA-sequencing data stem from mouse embryonic stem cells cultured in serum only while our cell population has been grown in the 2i and LIF medium. We conclude that Small-seq overall correlates well with bulk sequencing methods, within typical variability of distinct Omics methods and cell growth conditions.

### miR-183 and miR-294 are abundant and highly variable across single cells

miRNAs show highly variable expression between tissues (Keller et al. 2022), which is not surprising since cells with different tissue identity vary greatly at all layers of regulation from chromatin states to post-transcriptional regulation. However, little is known about the variability between individual cells belonging to the same tissues or cell culture. An advantage of single-cell sequencing over bulk sequencing is that variability of expression across cells can be measured. We first sorted the cells using principal component analysis (PCA), and as expected, our homogeneous cell population groups together without clear outlier cells (Figure 2A). However, while some cell pairs show remarkably similar expression profiles (Figure 2B) other pairs show substantial differences (Figure 2C). We next compared the mean expression of individual miRNAs against their expression variation across all cells (Figure 2D). As is typical of single-cell sequencing data, the mean expression and expression variation are inversely correlated – resulting in a negative slope - because of technical sampling noise (Grun et al. 2014). However, some miRNAs were clearly more variable than expected for technical reasons, indicating biological variability (miRNAs indicated in red, Figure 2D). Some miRNAs - exemplified by miR-16 - were confined to narrow ranges of expression while others - such as miR-183 and miR-294 - were variably expressed across individual cells (Figure 2E). We next compared the mean miRNA expression against miRNA half-lives, as measured in a previous study (Kingston and Bartel 2019). We found that highly variably miRNAs tend to have high expression and long half-lives (Figure 2F, top), while stably expressed miRNAs often have low expression and short half-lives (Figure 2F, below). We would expect that miRNAs with short half-lives are stable, since fast turn-over allows for rapid transcription, which in turn reduce expression variation (noise). It is surprising that highly expressed miRNAs tend to be more variable, but this could be specific to mESCs, where cell transitions including differentiation might depend on induction of a few specific miRNAs. Overall, we find that most miRNAs are stably expressed in mouse embryonic stem cells, while a few such as miR-183 and miR-294 are highly variable.

**Figure 2:**
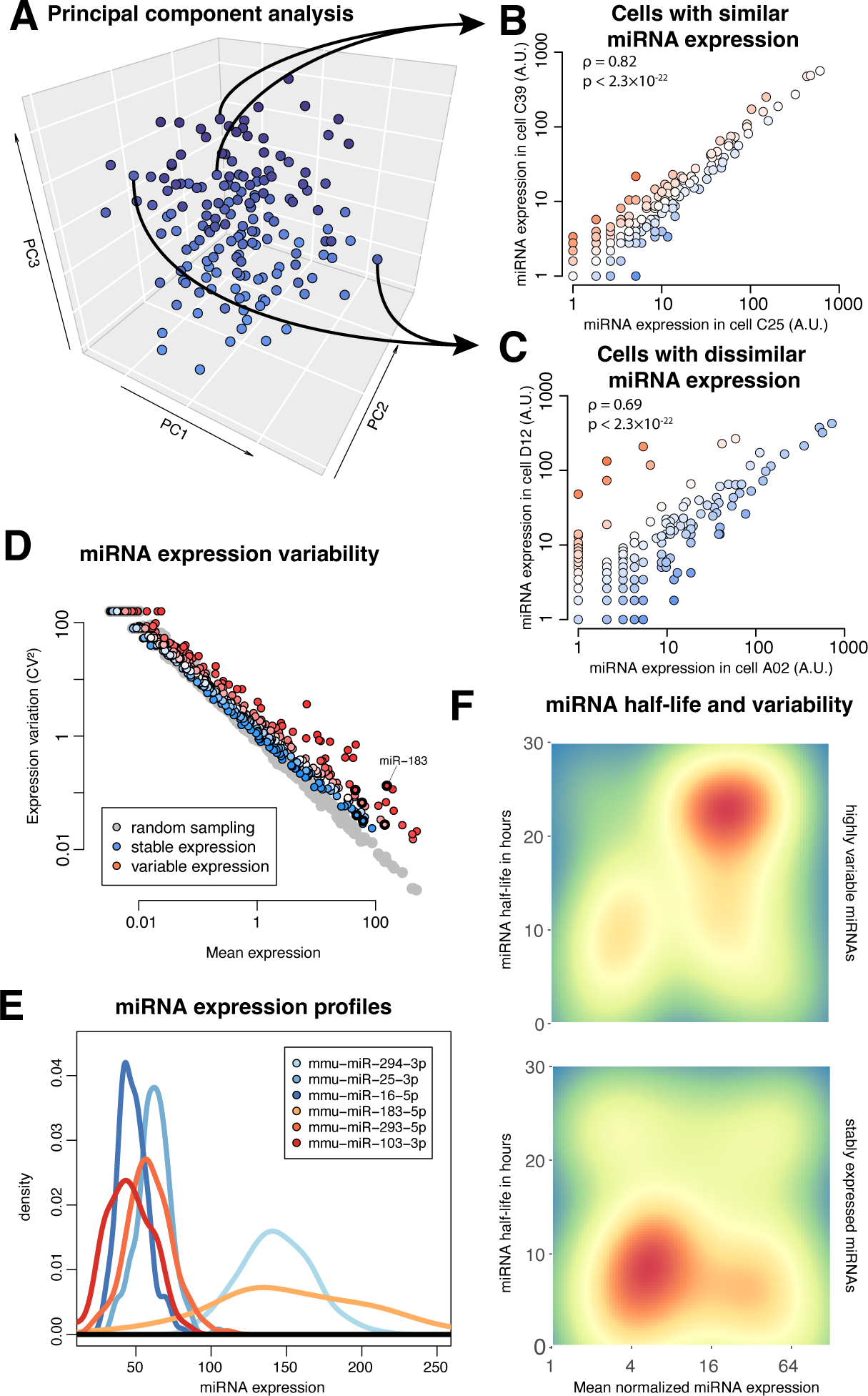
miRNA expression variation across single mouse embryonic stem cells. **A** Principal component analysis (PCA) of 192 single cells by their miRNA expression profiles. **(B)** Comparison of miRNA expression between two cells with similar profiles. Each dot indicates expression of one miRNA. **(C)** Comparison of miRNA expression between two cells with dissimilar profiles. **(D)** miRNA mean expression vs. expression variation. Each dot indicates one miRNA. Red color indicates variably expressed miRNAs while blue indicates stable expressed miRNAs. The grey dots are generated by random sampling and indicate expected technical noise. **(E)** Expression of six select variable (red) and stable (blue) miRNAs across 192 cells. The density profiles are smoothened. **(F)** Heatmaps of variably expressed (top) and stably expressed (bottom) miRNAs, indicating mean expression vs. miRNA half-life. Red color indicates the presence of several miRNAs with those features; blue color indicates the absence of miRNAs with those features.

### miRNAs form four distinct co-expression groups across single cells

Another advantage of single-cell sequencing is that covariations of miRNAs across single cells can be detected. Sorting miRNAs and cells according to their expression revealed clear global patterns (Figure 3A). We find several miRNAs that group together according to their expression profiles, including from top to bottom: (i) miR-26a, miR-30e, miR-182, miR-183 and miR-298; (ii) miRNAs that derive mainly from the 5-prime arms of the mir-290 genomic cluster; (iii) miRNAs that derive mainly from the 3-prime arms of the mir-290 cluster and (iv) a cluster comprised of miR-16, miR-25, miR-92, miR-103 and miR-130a. These expression trends are also observable when individual miRNAs are grouped into families based on their respective seed sequences (not shown). Curiously, there is a group of cells that express many mature miRNAs from both the 3-prime arm and the 5-prime arm of the miRNA genes from the mir-290 cluster (cell group ‘A’) and a group that express few mature miRNAs from both arms of genes from this cluster (group ‘C’). At the same time, there are other groups of cells which show discordant expression of these miRNA groups. Specifically, in group B, miRNAs stemming from the 3-prime arm of the mir-290 cluster are more abundant than those originating from the 5-prime arm, while in group E the reverse holds. This suggests that there may be unknown global protein cofactors present in individual cells that drive preferential processing of either arm of the hairpin (see below). Surprisingly, miR-293 seems to act contrary to all other members of the mir-290 cluster, with its 3-prime arm following the expression pattern of the 5-prime arm of the other miR-290 members, and its 5-prime arm following the pattern of the 3-prime arms of the other cluster members. In summary, we find that clustered miRNAs, but also some un-clustered miRNAs, follow each other in expression patterns, and that strand selection of miR-293 appears to be contrary to that of the other cluster members. These observations prompted a more in-depth analysis into the expression of individual miRNAs and global processing patterns.

**Figure 3:**
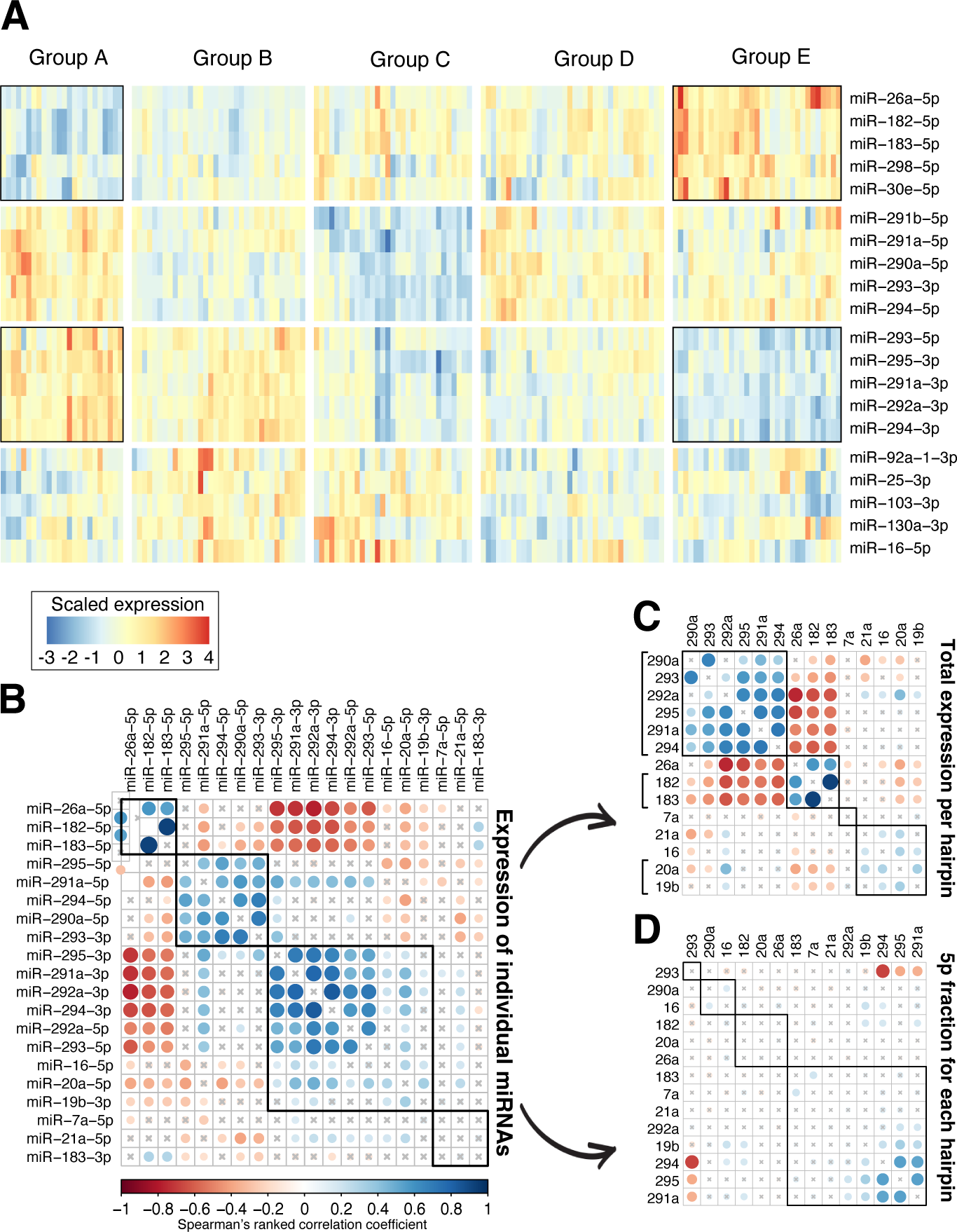
miRNA expression covariation across single mouse embryonic stem cells. **A** Four distinct miRNA co-expression groups across 192 cells. The miRNAs (rows) and cells (columns) were grouped using unsupervised hierarchical clustering. Red color indicates high expression relative to the mean for the given miRNAs, blue color indicates low expression. **(B)** Correlation between expression of miRNA strands. Blue color indicates positive correlation while red indicates negative correlation. **(C)** As in **(B)**, but summing expression from 5-prime and 3-prime arms of each miRNA hairpin. **(D)** As in **(B)**, but the correlation was done on the fraction of sequenced RNAs that originate from the 5-prime arm of the hairpin. Blue color indicates that the two miRNA hairpins have similar arm select across the cells, while red color indicates that the two hairpins have opposite arm selection across cells.

### miRNAs from the miR-182 and miR-290 clusters are anti-correlated in expression

We next changed our focus from expression patterns of groups of miRNAs to specific expression covariations of pairs of individual miRNAs (Methods). These analyses largely recapitulate three of the four groups identified above - (i) miR-26a, miR-182 and miR-183; (ii) miRNAs that derive mainly from the 5-prime arms of the mir-290 genomic cluster and (iii) miRNAs that derive mainly from the 3-prime arms of the mir-290 cluster but also including miR-16 and miR-20a (Figure 3B). The former is anti-correlated with the latter two, more so with the 3-prime arms than with the 5-prime arms. The two arms of the mir-290 cluster group more strongly among themselves but show no anti-correlation with one another. Summing up mature miRNAs from both arms to estimate overall miRNA precursor expression, we now see that all members of the mir-290 cluster group strongly together and are negatively correlated with miR-26a, miR-182 and miR-183 (Figure 3C). We can also clearly see the co-expression within the mir-182 and miR-183 genomic cluster, and within the mir-17 cluster (miR-19b and miR-20a). The observed covariation of miRNA originating from the same genomic clusters suggests that co-transcription is a major player in shaping miRNA expression profiles.

### Each single mouse embryonic stem cell has biases in miRNA arm selection

We next studied arm biases of miRNA precursors, by correlating the fraction of sequenced miRNAs that originate from the 5-prime arms of precursors, across cells (Figure 3D). It appeared that arm selection is not highly correlated between miRNA precursors, however, we can clearly see that the 5-prime fraction of miR-293 is strongly anti-correlated with that of other members of the mir-290 cluster, while these other members show positive correlations of their 5-prime fractions. Further, close observation reveals that in fact the 5-prime fractions of many miRNAs are subtly and positively correlated with each other and anti-correlated with the 5-prime fraction of miR-293 (Suppl. Fig. 3-4). This suggests that there may be global arm biases in certain cells, possibly mediated through the activity of a yet unknown protein cofactor that impacts all miRNAs equally - with the notable exception of miR-293.

### Pri-miRNAs from the miR-17 and miR-290 clusters are positively correlated in expression

While Small-seq is optimized for detecting mature miRNAs and does not detect miRNA primary transcripts (pri-miRNAs), we detect these in our previous study, where we applied the Smart-Seq2 method to profile poly-adenylated transcripts in single cells (Tarbier et al. 2020). In cells genetically depleted for Drosha we detect large parts of the pri-miRNA transcripts, which is expected since this protein is needed for cleaving the primary transcripts to produce miRNAs (Figure 4A, below). In control cells that express Drosha protein, we mainly detect cleaved transcripts (Figure 4A, above). This is consistent with previous observations that cleaved pri-miRNAs remain in the chromatin for some time after processing (Conrad et al. 2014), and also pri-miRNAs have previously been reported in sequencing of single nuclei (Elias et al. 2023), where they are located. The paucity of full-length pri-miRNAs in the control cells further suggests that pri-miRNA are cleaved relatively fast after being transcribed in mouse embryonic stem cells. We used the levels of pri-miRNAs in the Drosha knock-out cells as a proxy for transcription (Figure 4B) and found that the miR-17 and miR-290 clusters were strongly covarying (correlation coefficient 0.6, Figure 4C). This suggests that the covariation between the miRNAs from the miR-17 cluster and the 3p-derived miRNAs from the miR-290 cluster (Figure 3B) is induced at the transcriptional level, while the negative covariation between the miR-17 cluster and the 5p-derived miRNAs is induced post-transcriptionally, by an unknown mechanism. Unfortunately, we could not robustly profile expression of the mir-182 cluster in these data, so we cannot resolve if the anticorrelation between the mi-290 and mir-182 clusters are induced at the transcriptional level. Overall, we use single-cell data to infer expression of pri-miRNAs and infer their dynamics and interdependencies.

**Figure 4:**
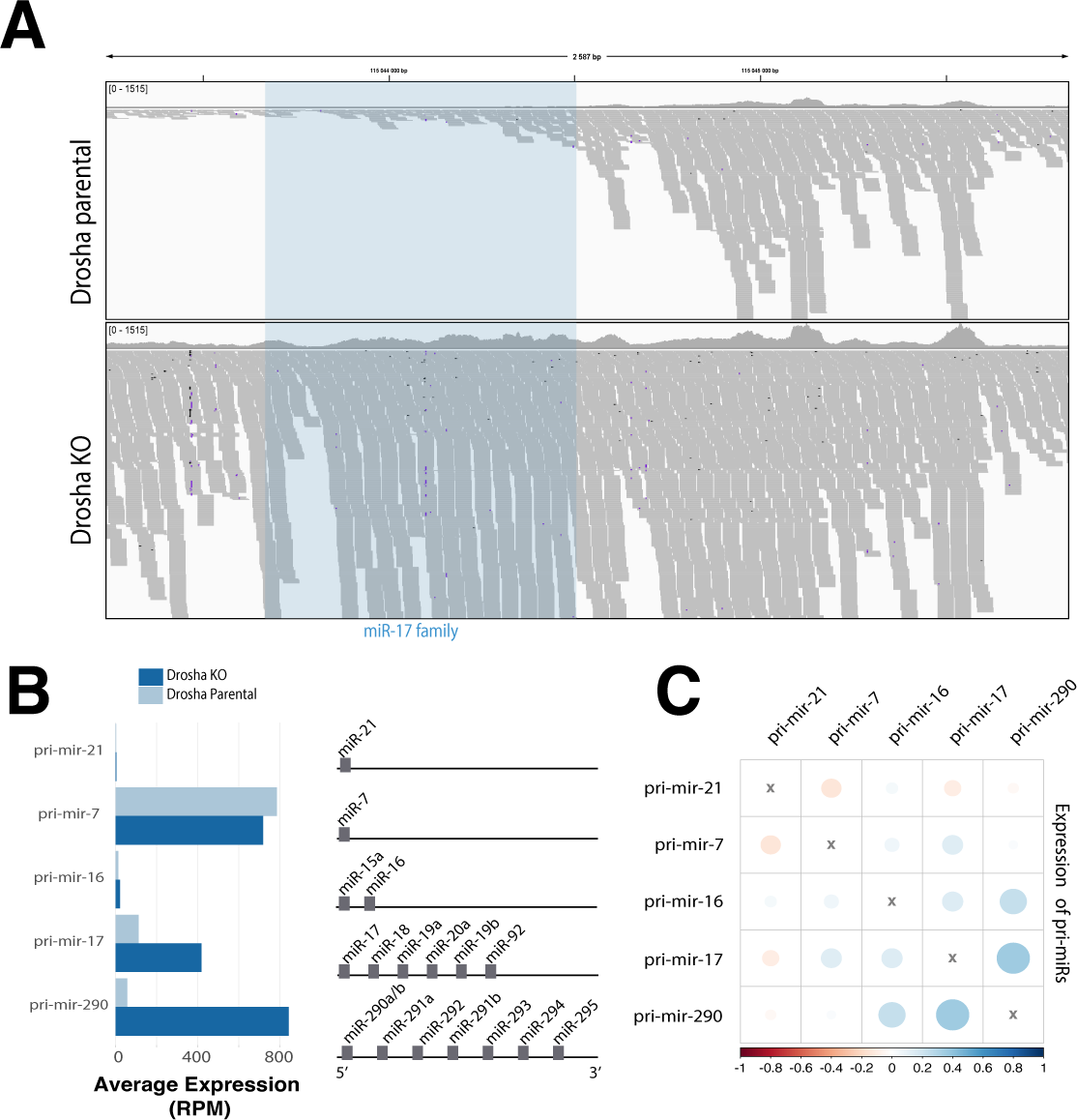
Expression covariation of miRNA primary transcripts. **A** Genome browser shot of miRNA primary transcripts from the miR-17 cluster sequenced by Smart-seq2. The clustered miRNAs (in blue shading) are cleaved out in the control cells (above) but retained in the Drosha KO cells (below). Each grey shading indicates one sequence read; the 3’ end of the primary miRNA is to the right. **(B)** miRNA primary transcript expression in control and Drosha KO cells (left). Schematic of mature miRNAs that are part of the same miRNA primary transcript (transcribed from the same genomic cluster). **(C)** Expression covariation of miRNA primary transcripts. The color code indicates the Spearman’s ranked test correlation value. Blue color indicates positively covarying primary transcripts.

### Most miRNA repression is conferred by miR-17, miR-291 and miR-292

We next investigated the impact of miRNAs on their target expression, by comparing single-cell RNA-seq data from Drosha knockout cells with control cells (Tarbier et al. 2020). We found that computationally predicted TargetScan targets (Agarwal et al. 2015) of the ten most highly expressed miRNAs were on average de-repressed around 15%, consistent with findings from bulk studies (Methods) (Baek et al. 2008; Selbach et al. 2008). However, this average repression effect was surprisingly small relative to the overall natural variation of expression between cells (Figure 5A), suggesting that the natural variation of expression overshadows the impact of the miRNAs at the RNA level (see Discussion section). We next looked at median changes of TargetScan targets in Drosha knockout versus control cells, for the top 16 miRNA families (Figure 5B-C). We found the miR-17, miR-291 and miR-292 families to have the strongest impact on the transcriptome, consistent with their relatively high measured expression in bulk and single cells (Figure 5B, panel inserts). The miR-290-5p and miR-291-5p families surprisingly have little impact on their targets’ expression, in spite of being fairly abundant. Similarly, it is surprising to see that the passenger strands of the miR-290 cluster are among the most abundant miRNAs but show practically no effect on their targets. This may suggest that these miRNAs are indeed just abundant by-products that are not loaded into Argonaute, or that they may have functions other than simple repression (see Discussion section below). To gain more confidence in these observations, we estimated mRNA half-lives transcriptome-wide through a time-series experiment of transcriptionally inhibited mouse embryonic stem cells. Specifically, we inhibited transcription with α-amanitin and measured mRNA levels at five time points following the transcriptional arrest and estimated half-lives using a linear model (Methods, Supplementary Table S2). We observed the same families to have strong target effects as in the previous analysis (Figure 5D), showing the robustness of these results and confirming that expression changes reflect changes in half-life rather than indirect effects at the transcriptional level. In summary, in our single cells we find that most repression is conferred by three families – miR-17, miR-291 and miR-292 – consistent with their high expression in bulk small RNA-seq data and also our mRNA half-life measurements.

**Figure 5:**
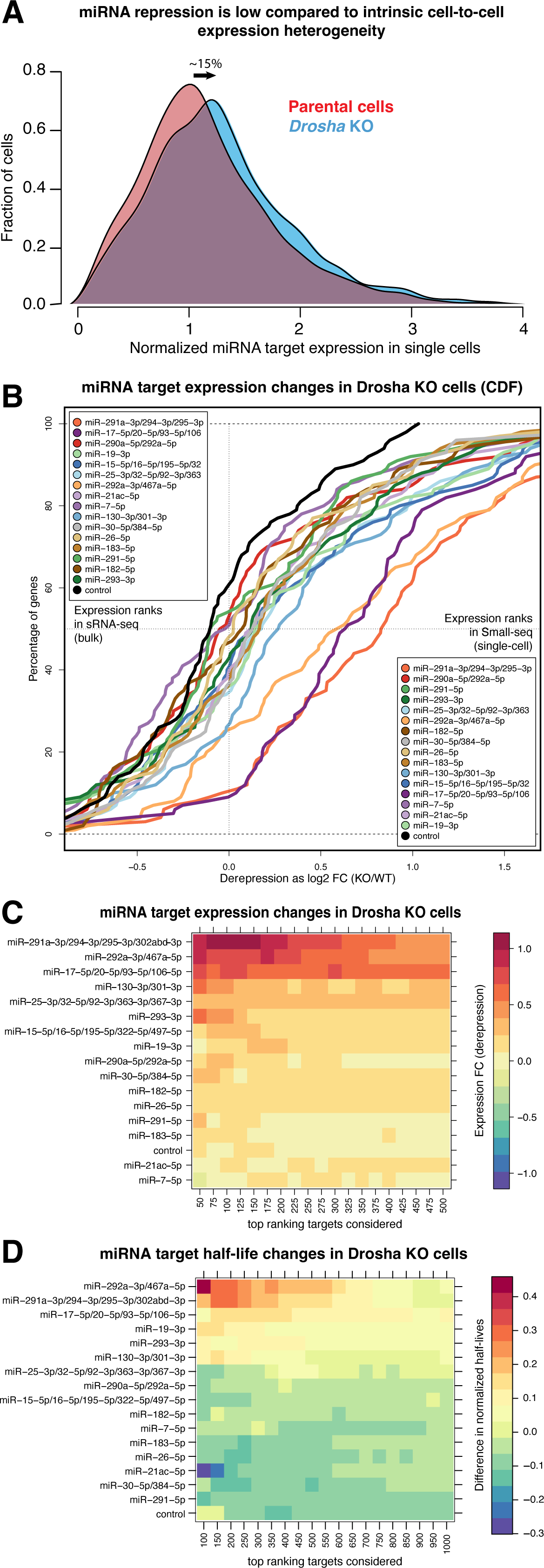
miRNA target repression in single cells. **A** Expression of miRNA targets in cells devoid of miRNAs (Drosha KO) and control cells (parental). The expression is normalized so that a cell with mean expression for the given target is assigned an expression value of 1. The density plot is made of a compound of the predicted TargetScan targets for the top 10 expressed miRNAs in mouse embryonic stem cells (Methods). The mean expression of the targets is upregulated (de-repressed) by ∼15% in the Drosha KO cells. **(B)** Cumulative distribution function (CDF) plots of the expression of targets of top miRNAs in mouse embryonic stem cells. The insert boxes show miRNA expression ranks according to bulk small RNA-seq (upper left corner) and Small-seq (lower right corner). **(C)** Heatmap of miRNA target de-repression in Drosha KO vs. control cells. The color indicates the log fold-change in expression, with red indicating stronger de-repression. The targets are sorted according to confidence level, as estimated by the TargetScan cumulative score. **(D)** as in **(C)**, but showing changes to transcript half-lives in Drosha KO vs. control cells.

### Distinct miRNAs increase and reduce target expression variation at RNA level

It is known from studies with reporter constructs, that miRNAs can buffer gene expression variation at the protein level (Schmiedel et al. 2015). Evidence from miRNA overexpression studies indicate that miRNAs may increase variation at the RNA level (Gambardella et al. 2017; Rzepiela et al. 2018), however, transcriptome-wide evidence from natural cell conditions is still lacking. We leveraged our published single-cell RNA-seq to and find that miRNA targets naturally tend to be more variable in expression at the RNA level than are other genes (Figure 6A). This particularly holds for targets of miR-291a, which is one of the most highly expressed miRNAs in mouse embryonic stem cells (Figure 5B). We further find that miRNA targets generally decrease in expression variation when Drosha is ablated, suggesting that they increase variation in their targets at the RNA level (Figure 6B). This in particular holds for targets of highly expressed and strongly repressing miRNAs like miR-291a, but also for miR-290a-5p, which although abundant, confers little repression (see above). Surprisingly, the targets of let-7 and miR-182 decrease in variability when Drosha is ablated, suggesting that these miRNAs naturally decrease variability of their targets at the RNA level. In summary, we provide evidence that miRNAs naturally induce expression variability of their targets at the RNA level, but surprisingly, select miRNAs have the exact opposite effect.

**Figure 6:**
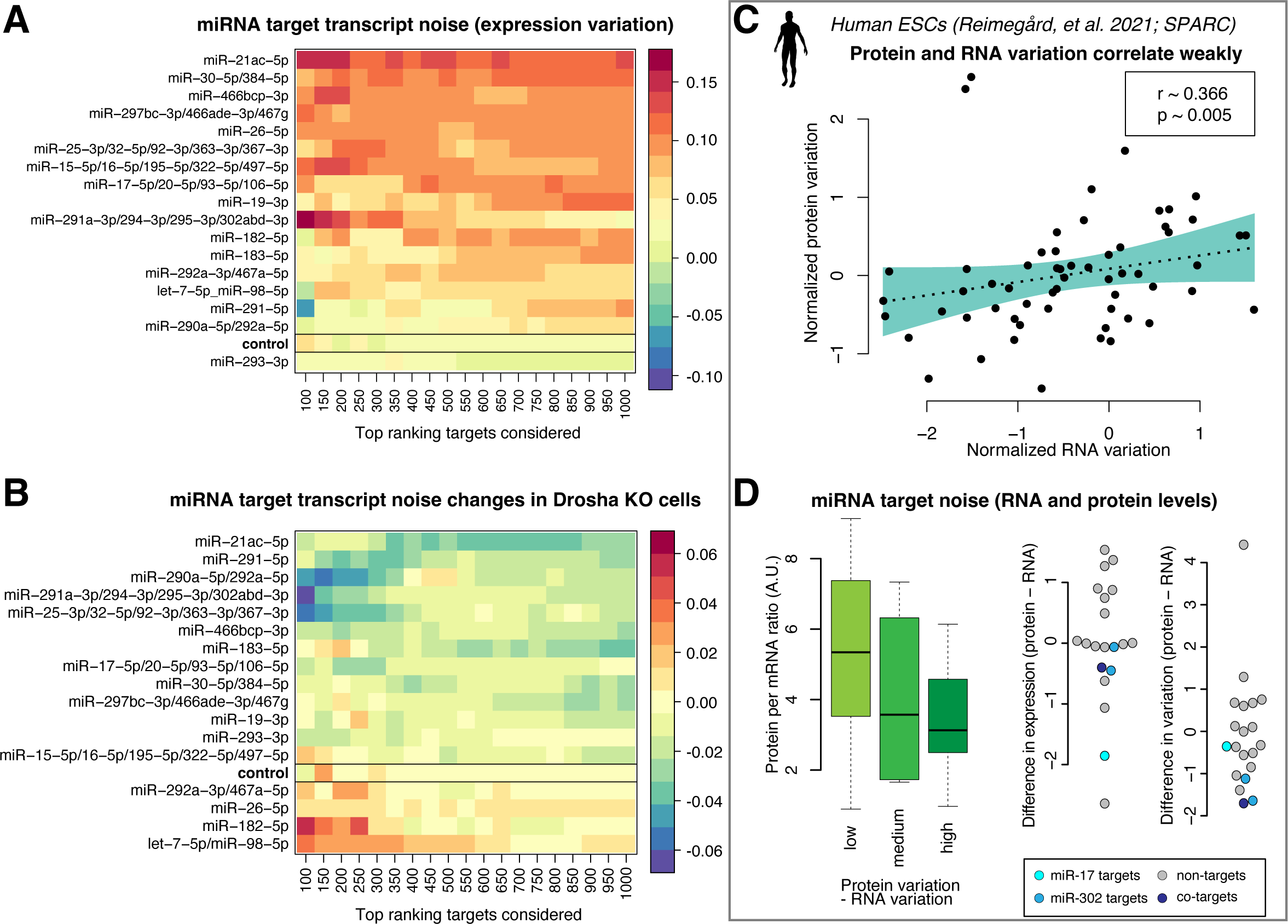
Expression variation of miRNA targets in mouse single embryonic stem cells. **A** Heatmap of miRNA target expression variation across single cells, without perturbation. The color code indicates expression variation (noise) as estimated by coefficient of variation squared (CV^2^) residuals (Methods). Red color indicates that the targets of the miRNA are naturally more variable. **(B)** As in **(A)**, but showing changes in expression variation following Drosha knockout. Blue color indicates miRNAs whose targets decrease in variation upon the loss of the Drosha biogenesis protein. **(C)** Expression variation for select genes at the RNA and protein level. Measurements were performed using combined single-cell RNA and protein profiling in the same single human embryonic stem cells (Reimegård 2021). Lower left panel: estimated translational efficiency and expression noise. The normalized RNA variation was for select genes subtracted from the normalized protein variation, and the genes were divided into three groups: genes with higher protein variation; genes with comparable RNA and protein variation; and genes with higher RNA variation. For each group, the estimated translational efficiency (Methods) was plotted. **(D)** difference between protein and RNA expression and difference between protein and RNA variation. Background genes were marked in grey, while miR-17 and miR-302 targets were marked in light blue and blue respectively. Genes that are regulated by both miRNAs are marked in dark blue.

### Evidence that targets of miR-17 and miR-302 are buffered at the protein level

We next investigated if RNA and protein variation correlates in single cells. In collaboration with colleagues, we recently developed SPARC - a new method to profile the whole poly-adenylated transcriptome and select proteins in the same single cells (Reimegard et al. 2021). Re-analyzing SPARC data from human embryonic stem cells, we found that RNA variation is a relatively poor predictor of protein variation for the same gene, in single cells (Figure 6C, above) (Reimegard et al. 2021). We found that proteins that are less variable than their cognate mRNAs tend to have high protein-to-mRNA ratios, while the reverse holds for proteins that are more variable (Figure 6D, lower left, Methods). Further, when estimating the ratio of RNA to protein, we found that the targets of miR-17 and miR-302, both abundant in human embryonic stem cells, are lower than for background genes (Figure 6D, lower right), suggesting translational repression. Last, we found that protein variation was lower than RNA variation for targets of miR-17 and miR-302, suggesting that the miRNA may increase RNA variation but buffer protein variation, consistent with previous studies on miRNA impact on expression noise (Figure 6D, lower right). However, these observations are preliminary and more methods development in the single-cell proteomics field will be required to elucidate this in a more systematic manner.

### Variable miRNAs induce strong gene expression covariations between target transcripts

It has previously been suggested that certain miRNAs may have the potential to induce covariations within their target pool (Rzepiela et al. 2018) and we have previously provided evidence that these effects may in fact occur naturally for many miRNAs (Tarbier et al. 2020). Here we re-analyze our single-cell RNA-seq data in more detail and find that targets of several specific miRNA families naturally covary in mouse embryonic stem cells (Figure 7A). Strikingly, most miRNA that induce such covariation patterns among their targets, such as miR-294-3p, were themselves highly variable (compare Figure 2E and 7A). Abundant miRNA families with low expression variation, such as miR-16 (part of the miR-15 family), do not seem to induce target co-expression (compare Fig. 2E and 7A). Surprisingly, miRNA abundance does not seem to determine covariation of the targets, rather the variability of the miRNA itself influences the level of covariations (Figure 7B). Since variation and covariation patterns within a cell sample cannot be measured with bulk methods, this again highlights the importance of single-cell miRNA quantifications in understanding miRNA function. A striking outlier in these analyses is the *let-7* family, whose targets are negatively covarying (asynchronous, Figure 7A). This miRNA family is lowly abundant in mouse embryonic stem cells but has been included for its well-described function in exit from pluripotency (Gambardella et al. 2017). Since it is lowly abundant in mouse embryonic stem cells and thus might not confer strong regulation of its targets, we speculate that its apparent effects in negative covariations could be linked to the function of its targets rather than direct miRNA effects. In summary, we provide evidence that several miRNA families can induce covariation between their targets, and that this particularly holds for miRNA families that are variable in expression between cells.

**Figure 7:**
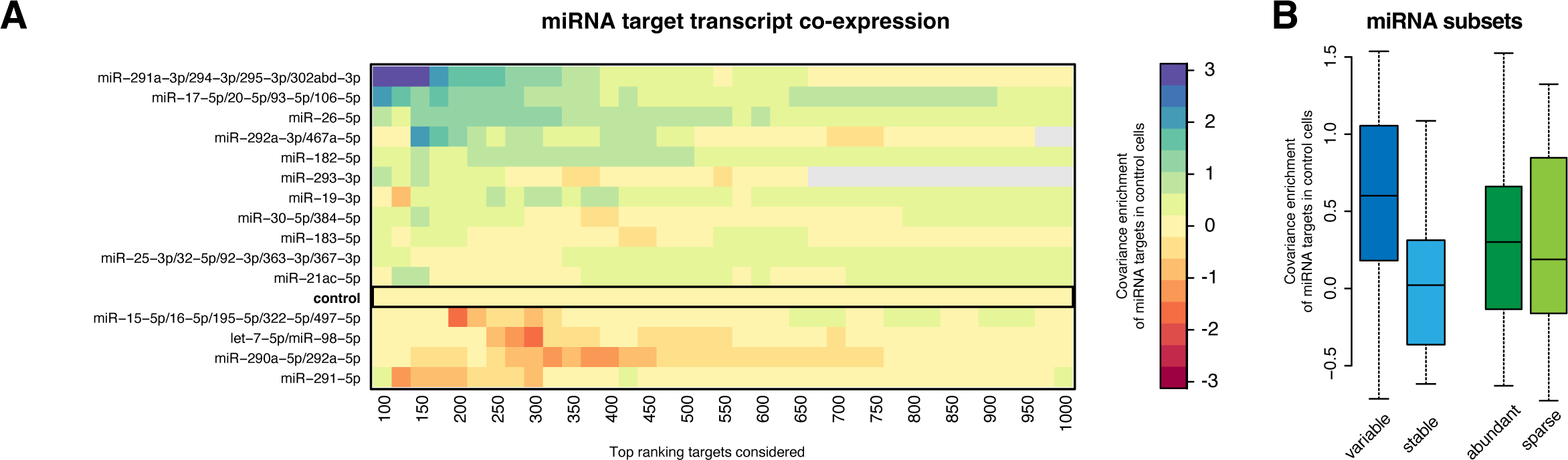
Expression covariation of miRNA targets in mouse single embryonic stem cells. **A** Heatmap of miRNA target covariation across single cells. The covariation enrichment was performed as previously described (Tarbier 2020). Blue color indicates mRNAs that are co-expressed more frequently than expected by chance; red color indicates mRNAs that are co-expressed less frequently than expected by chance. **(B)** Covariance enrichment for targets of miRNAs belonging to different groups, considering the top expressed 16 miRNA families. ‘Variable’ indicates the 8 miRNA families with the highest expression variation, while ‘stable’ indicates the 8 miRNA families with the most stable expression. ‘Abundant’ indicates the 8 top expressed miRNA families; while ‘sparse’ indicates the 8 least expressed miRNA families among this set.

## Discussion

Here we perform the first systematic and transcriptome-wide study of miRNA and target expression variation and covariation in single mouse embryonic stem cells. We show that miRNA profiles in single cells largely resemble their bulk counterparts although the single cell measurements can be used to estimate miRNA and target expression variation and covariation. We find that the sensitivity of miRNA single-cell sequencing is around 3%, which is lower than that of standard scRNA-seq. Most miRNAs are stably expressed in single cells, although a notable few are highly variable. In particular miRNAs from the mir-290 cluster and the mir-182 cluster are variably and negatively correlated. We show that miRNA profiles are shaped by the transcription of clusters and by a tendency for each single cell to favor miRNAs originating from either end of the hairpin precursors. This bias acts in a similar fashion on all miRNAs in a given cell and suggests the existence of unknown protein cofactors whose presence or absence drives this bias. Surprisingly, we find that the natural variation of mRNA expression appears to overshadow the repressive effect of miRNAs (∼15% repression), at least on the RNA level. Most miRNAs increase noise of their targets at the RNA level, but we provide evidence from published single-cell proteomics data that they may reduce noise at the protein level. Lastly, miRNAs that are themselves highly and variably expressed tend to induce expression covariations between their targets.

In the field of sequencing mRNAs, there exists methods that have sensitivity of >50% - meaning that probability that a given mRNA molecules will be represented in the generated sequence data (Hagemann-Jensen et al. 2020). In comparison, we here estimate that our method has a sensitivity of ∼3%, which is comparable or better than other methods tested (Hucker et al. 2021). This means that methods to sequence miRNAs in single cells are more dominated by stochastic sampling noise compared to ordinary scRNA-seq data, and sets limits to the quantitative questions that can be addressed. However, we here show that biological insights can be inferred from Small-seq data, when we focus on the 10-20 most highly abundant miRNA molecules. Limiting analyses to the most highly abundant miRNAs may not be a major problem, since the ten most expressed miRNAs both dominate the expression profile (Figure 1C) and also likely confer the most of the repression (Mullokandov et al. 2012).

We observe that the natural variation of expression of miRNA targets appear to overshadow the relatively subtle repression conferred by miRNAs, at least at the transcript level (Figure 5A). This could indicate that miRNAs may have important functions other than repression, such as regulating target expression variation or covariation. However, it should be considered that the target transcripts are translated to proteins, which typically have longer half-lives than transcripts. In this way the temporal fluctuations at the transcript level may be buffered at the more stable protein level. However, there is evidence for miRNA-induced noise reduction also at the protein level (Schmiedel et al. 2015), and for miRNA-induced covariation for proteins that form part of the same complexes (Gutierrez-Perez et al. 2021).

A main finding of our miRNA covariation analyses is that the mir-290 and mir-182 clusters are variably expressed and negatively correlated, suggesting a potential ‘switch-like’ mutually exclusive function between the two clusters. This is interesting, given that both clusters are involved in maintaining pluripotency and in stem cell self-renewal. Specifically, members of the mir-290 performs this function by indirectly upregulating Myc, Lin28 and Sall4 (Melton 2010), while the mir-182 cluster represses Argonaute2, a miRNA-interacting protein that can facilitate differentiation through for instance *let-7* (Suh 2011). In addition, members of the mir-290 cluster can advance progression through the cell cycle by targeting inhibitors of Cdk2-Cyclin E (Wang et al. 2008), and the mir-182 cluster can facilitate transitions between stem cell states by increasing variability of specific target genes such as Esrrb and Sox2 (Chakraborty 2020). It seems unlikely that differences between cells that have high mir-290 expression and cells with high mir-182 relate to cell cycle, given that the mouse embryonic stem cells in our study were sorted by cell stage (Methods). However, it seems possible that any switch-like behavior relates to different stem cell states, reflecting different compositions of pluripotency factors.

We further show that miR-293 is, in many ways, an outlier in the mir-290 cluster. It is the only abundant miRNA that targets GC-rich genes (Suppl. Fig. 7) and is an outlier with regard to its expression patterns and predicted targets in mouse embryonic stem cells (Ciaudo et al. 2009). miR-293 further appears to escape the global cellular precursor arm bias and in fact may be regulated exactly opposite from all other miRNA, including its polycistronic neighbors, by whatever factors drive said bias. This makes it an extremely interesting candidate for future research into miRNA processing.

Surprisingly, the passenger strands (5-prime arms) of the mir-290 cluster are among the most abundant miRNAs in mouse embryonic stem cells but have virtually no effect on their targets, as measured by mRNA steady-state levels and half-lives (Figure 5C-D). This suggests they can accumulate as by-products of the mature miR-290 miRNAs but are not loaded into Argonaute. Their stability has previously been confirmed through miRNA half-life estimates(Kingston and Bartel 2019).

Alternative miRNA functions are still understudied. The limited evidence we have to date stems from mathematical models, reporter assays, or overexpression experiments and is often limited to few miRNA-target interactions. We here reexamine existing hypotheses and add physiological evidence for thousands of miRNA-target interactions at the RNA level. Furthermore, we show that miRNA expression variability links to target covariation, highlighting the importance of inter-cellular miRNA heterogeneity.

In summary, while several studies have profiled miRNAs and their targets in single cells, there is a paucity of systematic and transcriptome-wide studies in individual mouse embryonic stem cells. We here outline the landscape of miRNA and target expression variation and covariation, and find that most miRNAs are stably expressed, with the notable exception a few miRNAs. The mir-290 and mir-182 clusters are variably expressed and negatively correlated, suggesting switch-like functions in defining distinct stem cell states. We further find that most miRNAs induce expression variation at the RNA level, but some may buffer variation at the protein level. miRNAs that are themselves highly and variably expressed induce covariations between their targets. Lastly, miRNA primary transcripts can be detected in whole-cell scRNA-seq data, opening up new possibilities in studying miRNA biogenesis at fine resolution.

